# Method to generate growth and degrowth models obtained from combinations of existing models applied to agrarian sciences

**DOI:** 10.1101/585224

**Authors:** André Luiz Pinto dos Santos, Guilherme Rocha Moreira, Frank Gomes-Silva, Cícero Carlos Ramos de Brito, Maria Lindomárcia Leonardo da Costa, Luiz Gustavo Ribeiro Pereira, Rogério Martins Maurício, José Augusto Gomes Azevêdo, José Marques Pereira, Alexandre Lima Ferreira, Moacyr Cunha Filho

## Abstract

Mathematical models that describe gas production are widely used to estimate the rumen degradation digestibility and kinetics. The present study presents a method to generate models by combining existing models and to propose the von Bertalanffy-Gompertz two-compartment model based on this method. The proposed model was compared with the logistic two-compartment one to indicate which best describes the kinetic curve of gas production through the semi-automated *in vitro* technique from different pinto peanut cultivars. The data came from an experiment grown and harvested at the Far South Animal Sciences station (Essul) in Itabela, BA, Brazil and gas production was read at 2, 4, 6, 8, 10, 12, 14, 17, 20, 24, 28, 32, 48, 72, and 96 h after the start of the *in vitro* fermentation process. The parameters were estimated by the least squares method using the iterative Gauss-Newton process in the software R version 3.4.1. The best model to describe gas accumulation was based on the adjusted coefficient of determination, residual mean squares, mean absolute deviation, Akaike information criterion, and Bayesian information criterion. The von Bertalanffy-Gompertz two-compartment model had the best fit to describe the cumulative gas production over time according to the methodology and conditions of the present study.

## 1. INTRODUCTION

Brazil has capacity and demand for the use of forage grasses as the main source of food in animal nutrition. However, the production capacity, nutritional value, and rumen degradation of the grass must be known to guide decisions to meet the nutritional needs of ruminants [12]. Diet formulation systems require knowing the nutritional value of foods, among which forage grasses. The kinetic parameters of degradation are important as they describe the digestion and characterize the intrinsic properties of foods that limit the availability to ruminants [17].

As reported by [5], several non-linear models are used to estimate the rumen fermentation kinetics of foods. A major advantage of those models is the possibility of biological interpretation of parameters [27]. However, when growth has a characteristic behavior that enables identifying steps, which allow dividing the curve into several stages, adopting multi-compartment models becomes necessary as they take exclusive parameters into account for each compartment [13].

A logistic two-compartment (LB) model was developed by [25] for kinetic studies of *in vitro* gas production based on the assumption that production rate is impacted by microbial mass and substrate level. Several researchers have used that model to study the kinetics of cumulative gas production [8,20]. However, the logistic model may not be adequate for some cases due to its fixed inflection point halfway through cumulative gas production [9]. [9,32] concluded that new models are still needed that can yield more biologically significant results with good mathematical fit of broad ranges of curve shapes with variable inflection points. In addition, creating new models for overall and specific situations is highly justifiable in face of the dynamics with which non-linear models have been applied in biological researches [2,23,24].

Thus, this study presents a method to generate growth and degrowth models by combining existing models and, specifically, to propose a new two-compartment model from the combination of the von Bertalanffy and Gompertz models. The logistic two-compartment model and the proposed one were compared to identify which has the best fit to cumulative gas production curves of ten genotypes of pinto peanut (*Arachis pintoi*) used in ruminant feed.

## 2. MATERIAL AND METHODS

### 2.1 Data Used

The genotypes were grown and harvested at the animal science station of CEPLAC in Itabela, BA, Brazil, a region located at 100 m altitude, 16°36’ S, and 39°30’ W featuring mean annual temperature of 23.3 °C and 1,350 mm of rainfall with no defined dry season. The genotypes were harvested in the rainier season. A randomized block experimental design with ten pinto peanut genotypes and three replicates was employed. The treatments comprised ten *Arachis pintoi* cultivars, namely: 13251 (G1), 15121 (G2), 15598 (G3), 30333 (G4), 31135 (G5), 31496 (G6), 31534 (G7), 31828 (G8), Itabela (G9), and Rio (G10). The genotypes were planted in beds with total area of 4 m^2^ and useful area of 1 m^2^. To obtain the dry matter (DM) and green matter (GM) production per hectare in both periods, the plants were cut 5 cm from the ground and, after the green forage was weighed, it was taken to the Animal Nutrition laboratory of the State University of Santa Cruz – UESC, where it was dried in a forced air circulation oven at mean temperature of 55 °C for 48 h and them ground in a Willey knife mill equipped with 1 mm sieve. The DM content at 105 °C was determined by drying until constant weight, crude protein (CP) and acid detergent insoluble protein (ADIP) were defined using the Kjeldahl method according to the AOAC [1], and neutral detergent fiber (NDF) and acid detergent fiber (ADF) were determined according to [31]. Gas production was read at 2, 4, 6, 8, 10, 12, 14, 17, 20, 24, 28, 32, 48, 72, and 96 h after the start of the *in vitro* fermentation process at the Federal University of Minas Gerais (UFMG) according to the equation proposed by [15].

### 2.2 Method to Generate Growth and Degrowth Models by Combining Existing Models

This section is one of the main objectives of our work. It consists in generalizing combination methods applied to agrarian sciences. These methods have been disseminated over several years in this area of science and, in the present work, in addition to gathering them, we provide other possible methods that, to the extent of our knowledge, have not been explored yet.

Let *W*_1_(*t*_1_,…,*t*_*k*_), …, *W*_*n*_(*t*_1_,…,*t*_*k*_) be existing models in the literature and consider 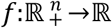 a function. Then

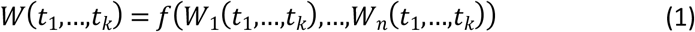

is a combination of such models via said function. Depending on the function, we can obtain several model-building methods, such as the ones below:

#### i) Method to generate growth and degrowth models via combinations in the weighted sums of power of models or linear combinations of power of existing models

Let *W*_1_(*t*_1_,…,*t*_*k*_), …, *Wn*(*t*_1_,…,*t*_*k*_) be existing models in the literature. Consider *f* 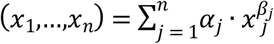, *x*_1_ = *W*_1_(*t*_1_,…,*t*_*k*_), …, *x*_*n*_ = *W*_*n*_(*t*_1_,…,*t*_*k*_), then:

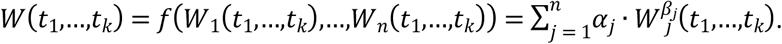

Therefore, for such function *f*, the building method is given by:

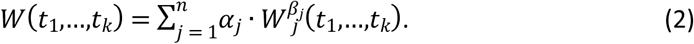

#### ii) Method to generate growth and degrowth models via combinations in the product of powers of existing models

In this case, use the function

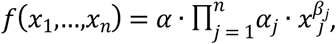

to obtain as building method

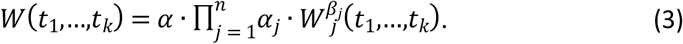

#### iii) Method to generate growth and degrowth models via combinations in the sum of products of existing models

In this case, simply use the function

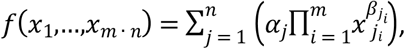

to obtain as building method

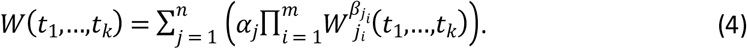

#### iv) Method to generate growth and degrowth models via combinations of the product of sums of existing models

In this case, we should consider the function

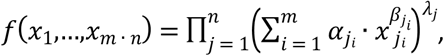

to obtain as building method

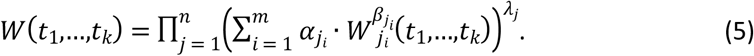

#### v) Method to generate growth and degrowth models via combinations in the sum of powers added to the product of powers of existing models

In this case, simply use the function

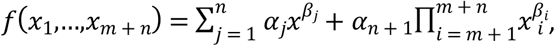

to obtain as building method

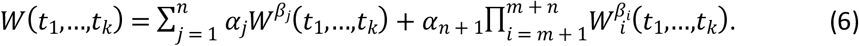

#### vi) Method to generate growth and degrowth models via combinations in the sum of powers of sums of existing models

The function stated as

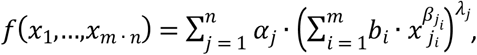

should be considered to obtain as building method

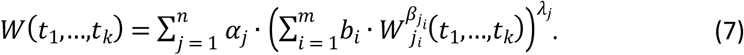

#### vii) Method to generate growth and degrowth models via combinations in adding parameters to existing models

Another building method occurs when parameters are added to an existing model. If *W*_1_ = *W*_1_(*t*_1_,…,*t*_*k*_/*β*_1_,…,*β*_*m*_), …, *W*_*n*_ = *W*_*n*_(*t*_1_,…,*t*_*k*_/*β*_1_,…,*β*_*m*_) are existing models, then for each function *f* and extra parameters *β*_*m* + 1_, … , *β*_*m* + *r*_, we can determine the building method:

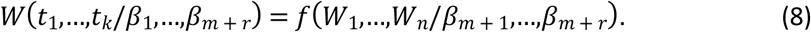

It can be seen that our method has a very broad character since Equation (1) may contemplate not only the time variable, but also other important variables for the comprehension of growth dynamics. Next, we present a “byproduct” of our method and illustrate it based on a numeric application.

### 2.3 Proposed Model and Theoretical Application

von Bertalanffy and Gompertz are basic models widely used to fit growth curves. The Gompertz model has been developed to describe microbial growth and was first used by [25] to study the kinetics of *in vitro* gas production [32]. These models were used in studies such as those by [16,30] for cumulative gas production kinetics. More recently, [24] presented the von Bertalanffy and Gompertz models, among others, (see Table 1, p. 2664) as sub-cases of what they called the method to generate growth models obtained from differential equations. Thus, the development of the mathematical model proposed resulted from the combination of the two models:

- 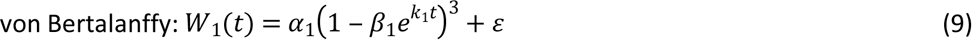
- 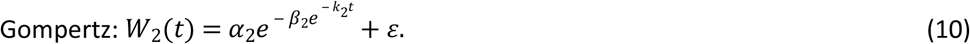

Indeed, let W(t) be an estimate of growth of the accumulated gas volume, hence, by building method (i) given by Equation (2), we can describe:

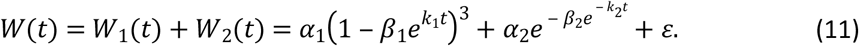

Thus, Equation (11) consists of our proposed model, called Two-Compartment von Bertalanffy-Gompertz model, or VGB, as it is a combination of Equations (9) and (10). Analogously, by observing the building methods provided in Section 2.2, we can generate one- or two-compartment models found in the literature, some of which fitted to *in vitro* gas production and others with potential application in this area, as described in Table 1.

**Table 1:** One- and two-compartment models from the methods

| Unicompartmental models | Building method | Models generated |
| --- | --- | --- |
| Brito-Silva [2] | (ii) | $W(t) = \alpha \left\{ (1 + \beta e^{-kt_f}) (1 + \beta e^{-kt_i})^{-1} e^{\lambda(\alpha^t_f - \alpha^t_i)} \right\}^\theta + \varepsilon$ |
| Exponential [16] | (vii) | $W(t) = \alpha_1 \{1 - e^{-k_1 t}\} + \lambda + \varepsilon$ |
| Logarithmic [16] | (vii) | $W(t) = \alpha / \{1 + e^{[2 + k(\lambda - t)]}\} + \varepsilon$ |
| Gompertz [16] | (vii) | $W(t) = \alpha e^{\{-e^{[1 - k(t - \lambda)]}\}} + \varepsilon$ |
| Two-compartment models | Building method | Models generated |
| Exponential-Logistic [33] | (i) | $W(t) = \alpha_1 \{1 - e^{(-k_1 t)}\} + \alpha_2 e^{\{-e^{[1 + k_2 \log(\lambda - t)]}\}} + \varepsilon$ |
| Michaelis-Menten [16] | (i) | $W(t) = (\alpha_1 t^c / t^c + k_1^c) + (\alpha_2 t^c / t^c + k_2^c) + \varepsilon$ |
| Gompertz Two-compartment [16] | (i) | $W(t) = \alpha_1 e^{\{-e^{(1 - k_1(t - \lambda))}\}} + \alpha_2 e^{\{-e^{(1 - k_2(t - \lambda))}\}} + \varepsilon$ |
| Exponential Two-compartment [19] | (i) | $W(t) = \{\alpha_1(1 - e^{-k_1 t}) + \lambda\} + \{\alpha_2(1 - e^{-k_2 t}) + \lambda\} + \varepsilon$ |
| Logistic Two-compartment [25] | (i) | $W(t) = \alpha_1 \{1 + e^{[2 - 4 \cdot k_1(t - \lambda)]}\}^{-1} + \alpha_2 \{1 + e^{[2 - 4 \cdot k_2(t - \lambda)]}\}^{-1} + \varepsilon$ |

In these models, *W*(*t*) is the accumulated volume (mL) at time *t*; *α* is the gas volume corresponding to complete substrate digestion (mL); *α*_1_ is the gas volume produced from the rapid-digestion fraction of non-fiber carbohydrates (NFC); α_2_ is the gas volume produced from the slow-digestion fraction of fiber carbohydrates (FC); *c*, *β*_1_, and *β*_2_ are shape parameters with no biological interpretation; *k* is the specific rate of gas production; *k*_1_ is the degradation rate of the rapid-digestion fraction (NFC); *k*_2_ is the degradation rate of the slow-digestion fraction (FC); *λ* is the time of bacteria colonization; *t* is the fermentation time; *e* is exponential; and *ε* is the random error associated with each observation with normal distribution, zero means, and constant variance. Thus, the cumulative gas production kinetics was fitted using models VGB and LB.

### 2.4 Estimating Parameters of Non-Linear Models, Assessors of Goodness-of-Fit, and Test of Model Identity and Parameter Equality

Next, the kinetics parameters of non-linear models VGB and LB were estimated via the least-squares method using the iterative Gauss Newton process through the function Nonlinear Least Squares. The statistical analyses were carried out using the software R version 3.4.1 [21].

To assess which model had the best fit, we used the following assessors: adjusted coefficient of determination 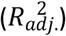, residual mean squares (RMS), mean absolute deviation (MAD), Akaike information criterion (AIC), and Bayesian information criterion (BIC) according to Table 2.

**Table 2:**
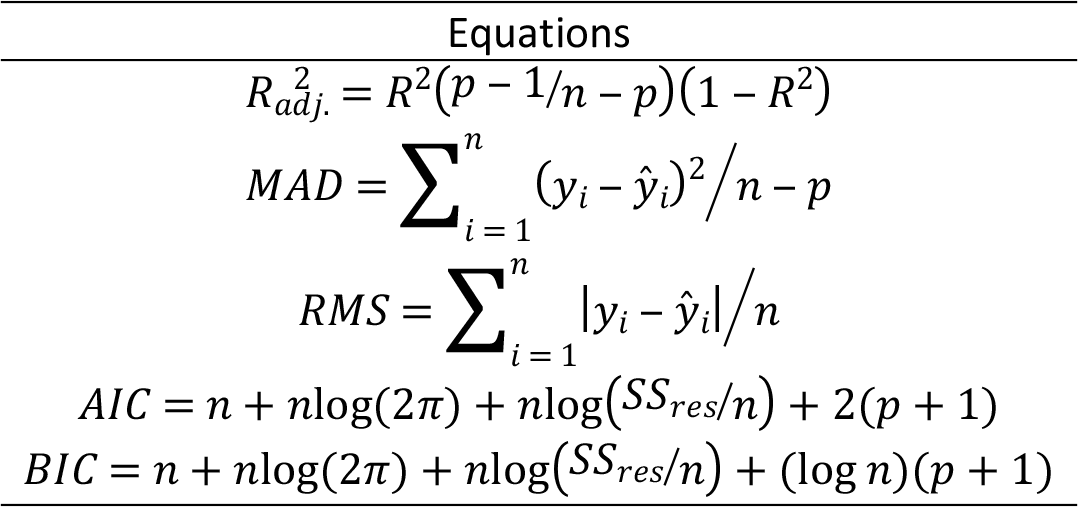
Mathematical description of the selection criteria

The terms that appear in Table 2 are described as follows: *SS*_*res*_ is the sum of the squares of the residues defined by 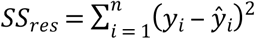, where *y*_*i*_ is the volume observed and 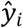 is the volume estimated (prediction) of *y*_*i*_; *n* is the number of observations, and *p* is the number of free parameters of the model. Thus, the best adjusted model is the one that has the lowest values for RMS, AIC, BIC, and MAD and the highest 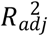 value.

## 3 RESULTS AND DISCUSSION

The cumulative gas production curves obtained from the observed and fitted data of genotypes of pinto peanut for both models had sigmoid shape over time and can be split into three stages, namely: initial stage of low gas production; exponential stage of rapid gas production; and asymptotically null stage or low gas production (Figure 1).

Models LB and VGB fitted to all stages of the fermentation process of genotypes G1, G2, G4, G5, G7, G8, G9, and G10. For genotypes G3 and G6, the models showed good fits both in the initial portion of the curve and in the exponential stage, but there is evidence they did not have good fits to the asymptotic phase, in which gas production was over- or underestimated, respectively, by the LB and VGB models. However, model behavior also largely depends on the morphological [4] and chemical [26] characteristics employed. The same model may have low or high performance when using genotypes of the same species or when using different substrates [30].

**Figure 1:**
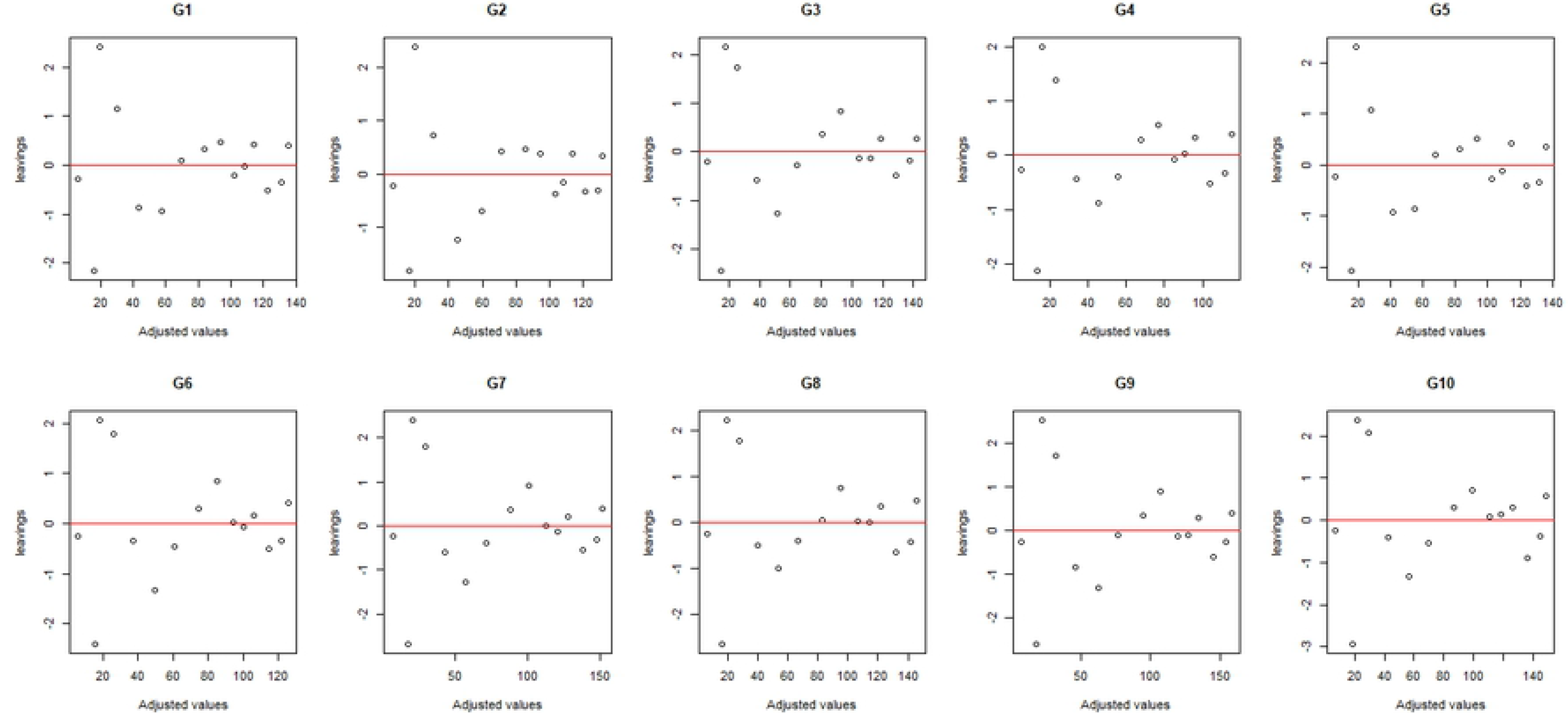
Cumulative gas production curves of the ten genotypes over incubation time based on the observed data and data fitted by models VGB and LB

Verifying the assumptions for the regression models is a very important step since, in case they are not met, the model is considered inadequate and such deviation must be corrected or taken into account in the model. Thus, in addition to verifying the goodness-of-fit by Figure 1, it is important to analyze the residues to verify the assumptions of the model. In order to asses goodness-of-fit through the analysis of residues, we can use the scatter plot of the residues as a function of the fitted values (Figure 2) and the quantile-quantile plot with the envelope of residues (Figure 3). The residue diagnostic plots provide no reason to deny the model assumptions have been met.

**Figure 2:**
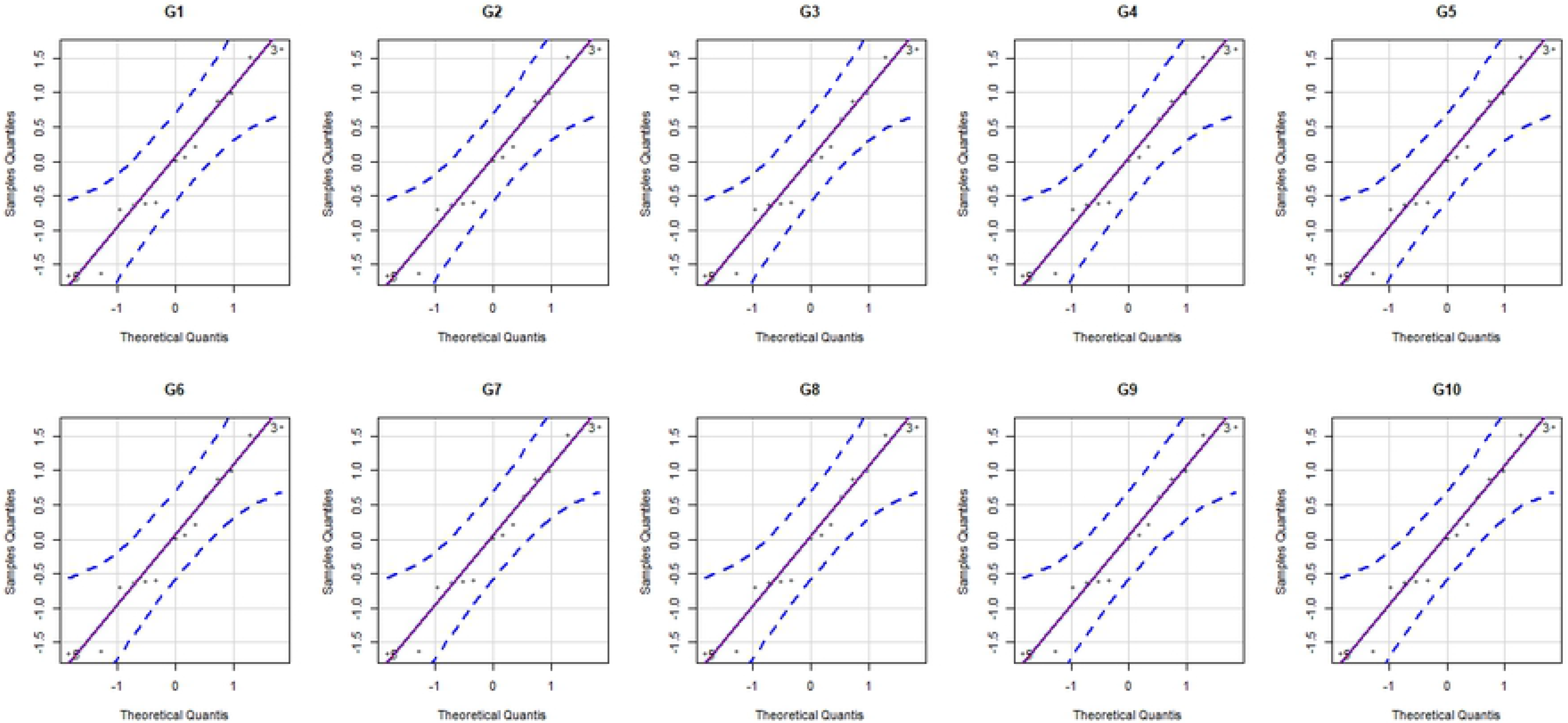
Scatter plot of the statistical model through the residues for all genotypes

**Figure 3:**
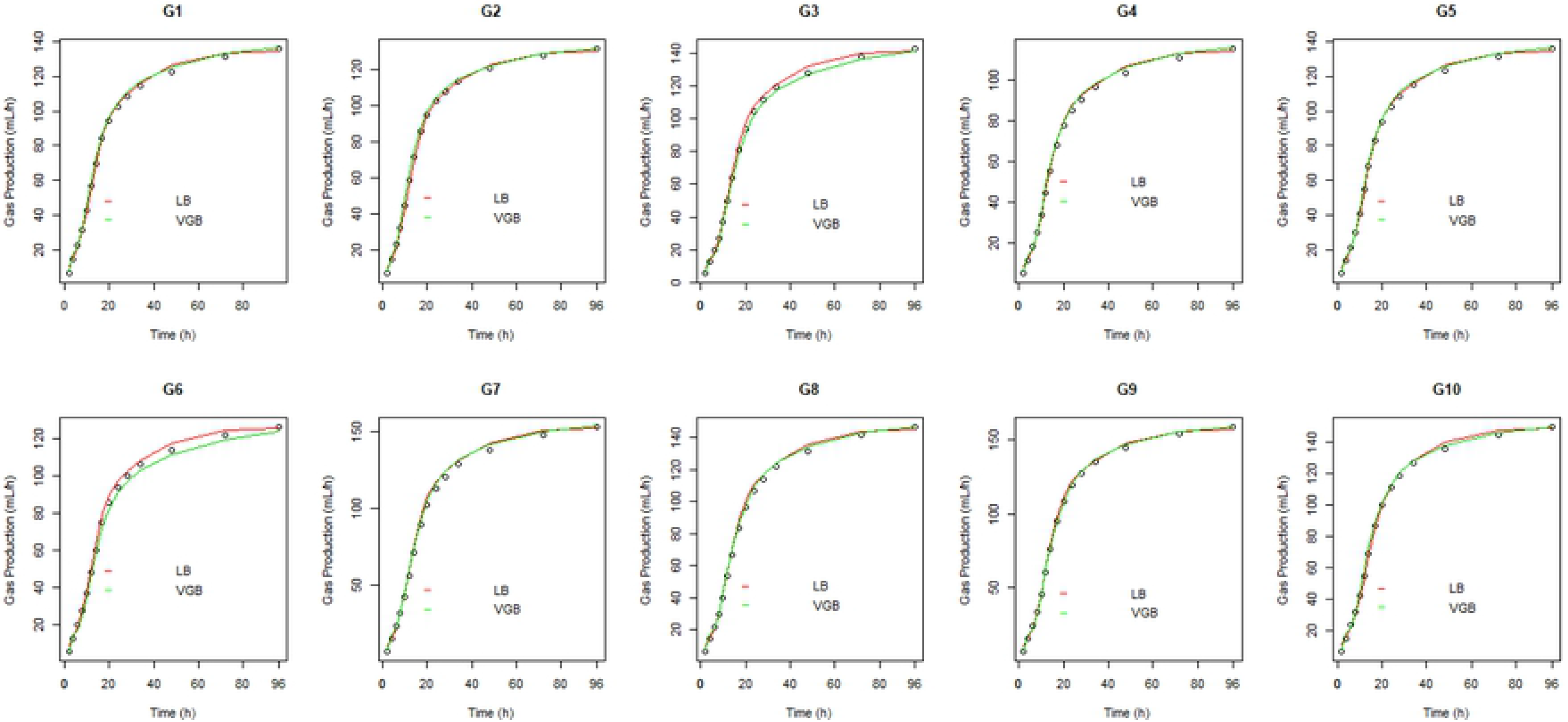
Normality plots of the statistical model through the residues for all genotypes

The models studied obtained 100% convergence and all kinetic parameters of degradation estimated by the different models were significant at 95% confidence. Colonization times (*λ*) ranged from 4.40 h for G2 to 5.46 h for G3. [6] fitted model LB to ten genotypes of *Arachis pintoi* and found similar *λ* values as those obtained in the present study at 4.4 to 5.5 h. Lower values were found by [7] for the *Arachis pintoi* cultivars assessed, from 2.8 to 4.3 h. Considering all genotypes, estimates 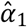 and 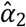 for model LB were higher and lower, respectively, than estimates 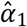 and 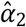 for model VGB. The final gas volume *W(t)* is produced by the rapid- and slow-digestion fractions, i.e., the sum of *NCF* and *FC*. Genotype G4 had the lowest total gas volume *W*(*t*) for models VGB and LB, whereas genotype G7 and G9 had the highest values for both models. According to [18], the volume of gases produced depends on substrate composition, i.e., the higher the starch and fiber contents, the lower and higher their gas productions, respectively.

According to Table 3, the estimated rates of rapid and slow degradation of the *NFC* and *FC* were 0.20 and 0.04 for the VGB model and 0.07 and 0.02 for the LB model, noting the dissimilarity among the genotypes for the two models evaluated with higher values for VGB. The estimated degradation rate values *k*_1_ and *k*_2_ of the VGB model were similar to those found by [3] for different forage grasses. Those authors reported values of 0.095 (0.04), 0.108 (0.04), 0.131 (0.04), 0.203 (0.04), 0.216 (0.04), and 0.222 (0.04) for silages of corn, alfalfa hay, sorghum, sugar cane, Coastcross hay, and Tifton-85, grass, respectively. The estimated degradation rates *k*_1_ and *k*_2_ by model LB were similar to those found by [6,7] for pinto peanut genotypes. Another relevant piece of information is that 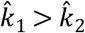 for the models fitted considering all genotypes (Table 3).

That matches the important aspect of the theory according to which parameter *k*_1_ is greater than parameter *k*_2_, i.e., *NFC* are more quickly degraded than *FC* [25,14]. [22] used this recommendation in their study.

**Table 3:**
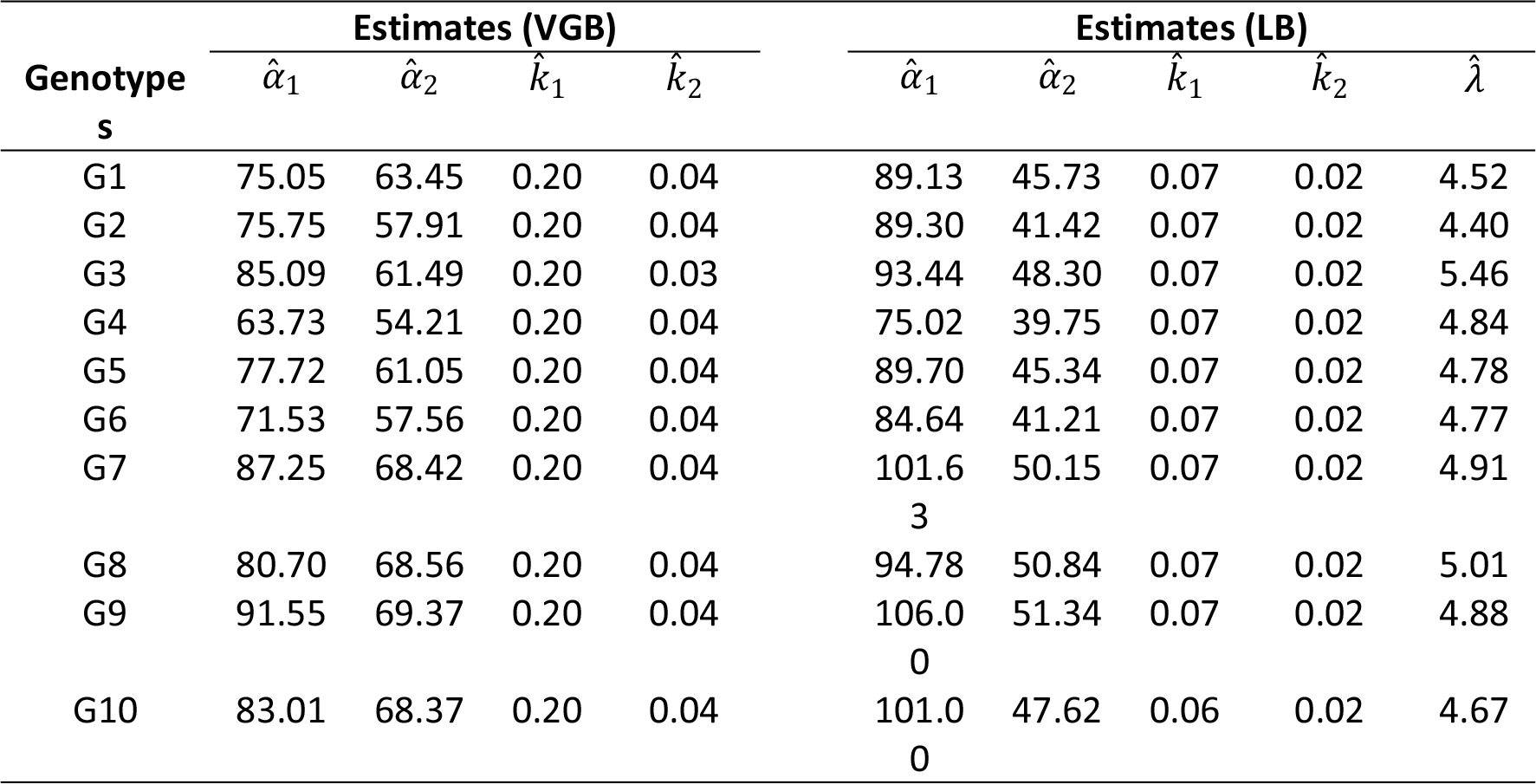
Estimated values of parameters *α*_1_, *α*_2_, *k*_1_, *k*_2_, and *λ* for the VGB and LB models fitted to data on pinto peanut genotypes

The goodness-of-fit assessors are presented in Table 4. Choosing the best models has not been an easy task since each of the different goodness-of-fit assessors proposed in the literature recommends a certain characteristic such as model simplicity [28]. However, the higher the number of assessors considered, the more adequate the indication of the best model(s) [29]. A comparison of the two models showed the smallest differences were found for 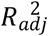, whose values were very close for both models, with no evidence of which has the best fit. Nevertheless, for all genotypes, when criteria RMS, MAD, AIC, and BIC were analyzed, we observed that the VGB model had the lowest values (Table 4). The best fitted model is the one that has the lowest values for RMS, AIC, BIC, and MAD and the highest 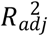 value. Therefore, the information favors indicating the best fits to the proposed model.

Few papers are found in the literature that reference studies on mathematical models for *in vitro* gas production using pinto peanut genotypes. [10], when comparing several models to assess pinto peanut genotypes during the rainier and less rainy seasons in Itabela, BA, Brazil, found the best fit through LB, followed by the von Bertalanffy, Gompertz, Brody, and Logistic models. Meanwhile, [16], when assessing the Brody, von Bertalanffy, Gompertz, France, logistic, modified logistic, and LB models to describe cumulative gas production in sunflower and corn silages, concluded the LB was the best model. According to [25,11], multi-compartment models had better goodness-of-fit than those based on first-order kinetics.

**Table 4:**
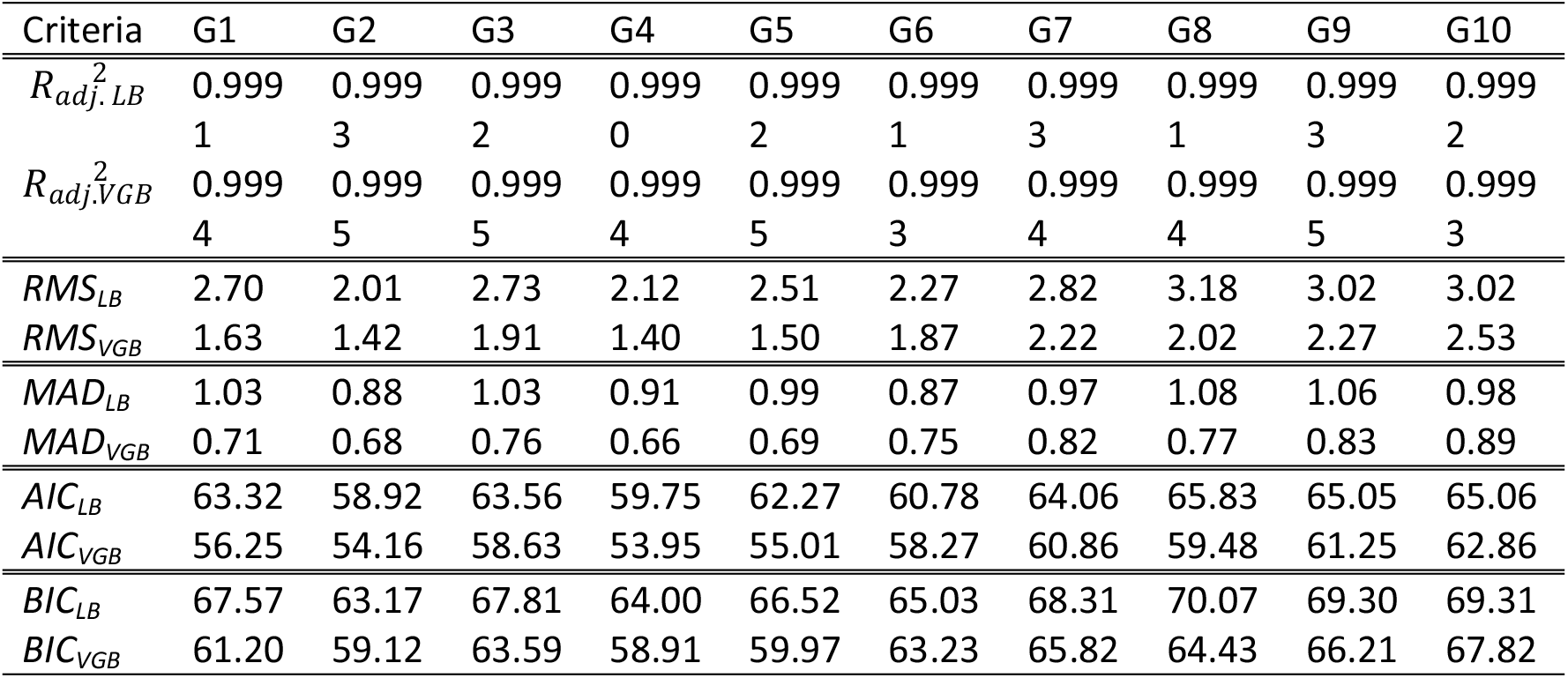
Criteria used to select the most adequate non-linear model

## 4 CONCLUSION

The results obtained indicated the VGB model may be an alternative to describe *in vitro* gas production curves. As it had good quality assessors and based on the biological interpretations of parameters, the VGB model proved superior to the LB one to describe growth curves. Therefore, it is recommended for the study of gas production kinetics from forage grasses in genetic enhancement programs according to the methodology and conditions under which the present study was developed.

